# Redox control of the deubiquitinating enzyme Ubp2 regulates translation during stress

**DOI:** 10.1101/2024.04.30.591912

**Authors:** Clara M. Santos, Blanche K. Cizubu, Dinachi Okonkwo, Chia-Yu Chen, Natori Maske, Nathan A. Snyder, Vanessa Simoes, Erica J. Washington, Gustavo M. Silva

## Abstract

Protein ubiquitination is essential to govern cell’s ability to cope with harmful environments by regulating many aspects of protein dynamics from synthesis to degradation. As important as the ubiquitination process, the reversal of ubiquitin chains mediated by deubiquitinating enzymes (DUBs) is critical for proper recovery from stress and re-establishment of proteostasis. Although it is known that ribosomes are decorated with K63-linked polyubiquitin (K63-ub) chains that control protein synthesis under stress, the mechanisms by which these ubiquitin chains are reversed and regulate proteostasis during stress recovery are still illusive. Here, we showed in budding yeast that the DUB Ubp2 is redox regulated during oxidative stress in a reversible manner, which determines the levels of K63-ub chains present on ribosomes. We also demonstrate that Ubp2 is a processive enzyme whose activity is modulated by a series of repeated domains and the formation of important disulfide bonds. By combining, cellular, biochemical, and proteomics analyses, we showed that Ubp2 is crucial for restoring translation after stress cessation, indicating an important role in determining cellular response to oxidative stress. Our work demonstrates a novel role for Ubp2, revealing that a range of signaling pathways can be controlled by redox regulation of DUB activity in eukaryotes, which in turn will define cellular states of health and diseases.

## Introduction

Eukaryotic cells are constantly exposed to everchanging environments, in which they must reprogram several physiological pathways to adapt and thrive. Under stress conditions, cells regulate gene expression at the transcription and translation levels, in addition to reshaping the functional proteome through post-translation modifications (PTMs), protein interactions, and degradation (1, 2). Several studies have focused on understanding a myriad of regulatory processes that occur upon stress induction (2–4), but the mechanisms by which cells recover from stress and re-establish proteostasis have remained more elusive. In response to oxidative stress, we observed in the budding yeast *Saccharomyces cerevisiae* that ribosomes are rapidly and heavily decorated with K63-linked polyubiquitin (K63-ub) chains (5–7), a less conventional type of polyubiquitin chain that functions independently of the proteasome (8, 9). We showed that the accumulation of these ubiquitin chains under oxidative stress is required to pause translation globally at the elongation stage (7, 10, 11), a process that is important to support cellular resistance to stress. We also determined that ribosome ubiquitination is promoted by the E2 ubiquitin conjugating enzyme Rad6, which is itself redox-regulated and whose activity is critical for reprogramming protein synthesis upon oxidative stress induction (5, 11). Although we have identified important genes and characterized new processes in the early stage of this pathway of Redox control of Translation by Ubiquitin (RTU), the mechanisms involved in the reversal of this process remain largely unknown.

Ubiquitination is highly dynamic process and during cellular recovery from oxidative stress, we have showed that K63-ub is rapidly reversed by the deubiquitinating enzyme (DUB) Ubp2 through a degradation-independent process (5). Ubp2 is a multifunctional DUB largely known for its role in regulating and antagonizing the E3 ubiquitin ligase Rsp5 in mediating trafficking of various types of membrane associated cargoes (12–14). Ubp2 is also known to participate in mitochondria homeostasis, protein quality control, and DNA damage response (15–19). Ubp2 is the largest DUB encoded in yeast (146 kDa), and even though it participates in several physiological processes, the structural and functional mechanisms regulating Ubp2 activity in the RTU and during the stress response are not understood.

DUBs are proteases that cleave the isopeptide bond between the C-terminus glycine of ubiquitin and the ε-NH_2_ group of a lysine residue in a substrate or ubiquitin itself when in the form of a chain. DUBs can then trim, remove, or even remodel selective ubiquitin chains (20–22). In addition, DUBs are functionally classified based on their catalytic activity, which are divided into cysteine proteases or metalloproteases (20, 23, 24). Cysteine proteases are far more abundant and because of their functional and structural diversity, several mechanisms of regulation have been identified via PTMs, allosteric regulation, protein trafficking, and /or protein-protein interaction (20). In addition, because of their catalytic cysteine, DUBs can be redox-regulated by participating in a series of oxidation and reduction reactions that reversibly modulate their activity (20, 25, 26). A previous study has shown that the activity of DUBs can be either unaffected, enhanced, or activated by the thiol reducing agent DTT (dithiothreitol), suggesting that DUB’s protein sequences and structural features can determine their sensitivity to reactive oxygen species (ROS) (27). However, the nature of their regulation and the physiological impact of these redox modifications remains largely unknown. As a cysteine protease from the USP/UBP family, redox regulation of Ubp2 can be critical to regulate proteostasis during cellular stress.

Here, we show that the activity of the cysteine DUB Ubp2 is inhibited by hydrogen peroxide (H_2_O_2_) and reversibly regulated during the oxidative stress recovery. Furthermore, reactivation of Ubp2 is required to reverse K63-ub chains and its activity is controlled by selective repeated domains and reactive cysteine residues. Finally, we showed that Ubp2 is critical for the re-establishment of proteostasis supporting translation resumption after stress cessation. Our work provides new insights on the redox regulation of Ubp2 and proposes new models by which several ubiquitin mediated pathways could be rapidly induced by common and prominent stressors, with meaningful impact to cellular physiology and health.

## Results

### Ubp2 is a linkage-specific DUB regulated by ROS

We previously identified that deletion of *UBP2* impairs cellular capacity to reverse K63-ub chains that accumulate in response to oxidative stress in yeast (5). However, it remained unknown whether Ubp2 is directly responsible for the cleavage of K63-ub chains as part of the RTU and whether Ubp2 activity could be regulated by ROS. Expression of wild-type (WT) Ubp2 in *ubp2*Δ rescues the deubiquitination phenotype, which reverses K63-ub globally but also from ribosomes (Figs. 1*A*, S1). By mutating Ubp2’s catalytic cysteine residue to a serine (C745S), we confirmed that Ubp2 activity is required for the reversal of K63-ub from ribosomes that accumulate under H_2_O_2_ (Fig. 1*A*). To further test Ubp2’s role in directly cleaving these K63-ub chains, we purified recombinant Ubp2 and showed that Ubp2 acts preferentially on K63-ub in comparison to K48-polyubiquiting linked chains (K48-ub) (Fig. 1*B*). We also showed that Ubp2 is a processive enzyme able to generate degradation intermediates of varied lengths when tested against synthetic tetra- and hexa-K63-ub chains (Fig. 1*C*). Combined, our results suggest that Ubp2 is able to specifically trim K63-ub directly from its targets. Given that Ubp2 is a cysteine protease, we hypothesized that the accumulation of K63-ub under oxidative stress occurs due to oxidative inactivation of Ubp2. To test this possibility, we performed *in vitro* activity assays, and we observed that the catalytic activity of Ubp2 against the fluorogenic substrate Ub-AMC can be inhibited by H_2_O_2_ in a dose-dependent manner (Figs. 1*D*, S2, *A and B*). As expected, the pan DUB inhibitor PR-619 and the cysteine alkylator iodoacetamide (IAA) abrogated Ubp2 enzymatic activity. Importantly, oxidative reduction of Ubp2 activity can be rescued by the reducing agent dithiothreitol (DTT), which indicates a reversible regulatory process (Fig. 1*D*). These results suggest that H_2_O_2_ promotes the oxidation of Ubp2’s catalytic cysteine residue, which can be reversed by thiol reducing agents. This redox regulation also occurs when Ubp2 is tested against synthetic K63-ub chains (Fig. 1*E*). While we observed a redox regulation of recombinant Ubp2 *in vitro*, it was unclear whether this process also occurred in the cellular context. To test that, we immunoprecipitated Ubp2 from yeast cells subjected to oxidative stress and we showed that Ubp2’s activity is partially inhibited by H_2_O_2_, which can also be rescued by DTT (Fig. 1*F*). HA-immunoprecipitation from cells expressing the catalytic dead Ubp2^C745S^ did not display activity against Ub-AMC, demonstrating that this deubiquitinating activity was produced by Ubp2 and not by other contaminant proteases (Fig. 1*F*). Therefore, our results indicate that Ubp2 activity is redox-regulated by H_2_O_2 and_ Ubp2 is responsible for the reversal of K63-ub chains that accumulate on ribosomes under stress.

**Figure 1.**
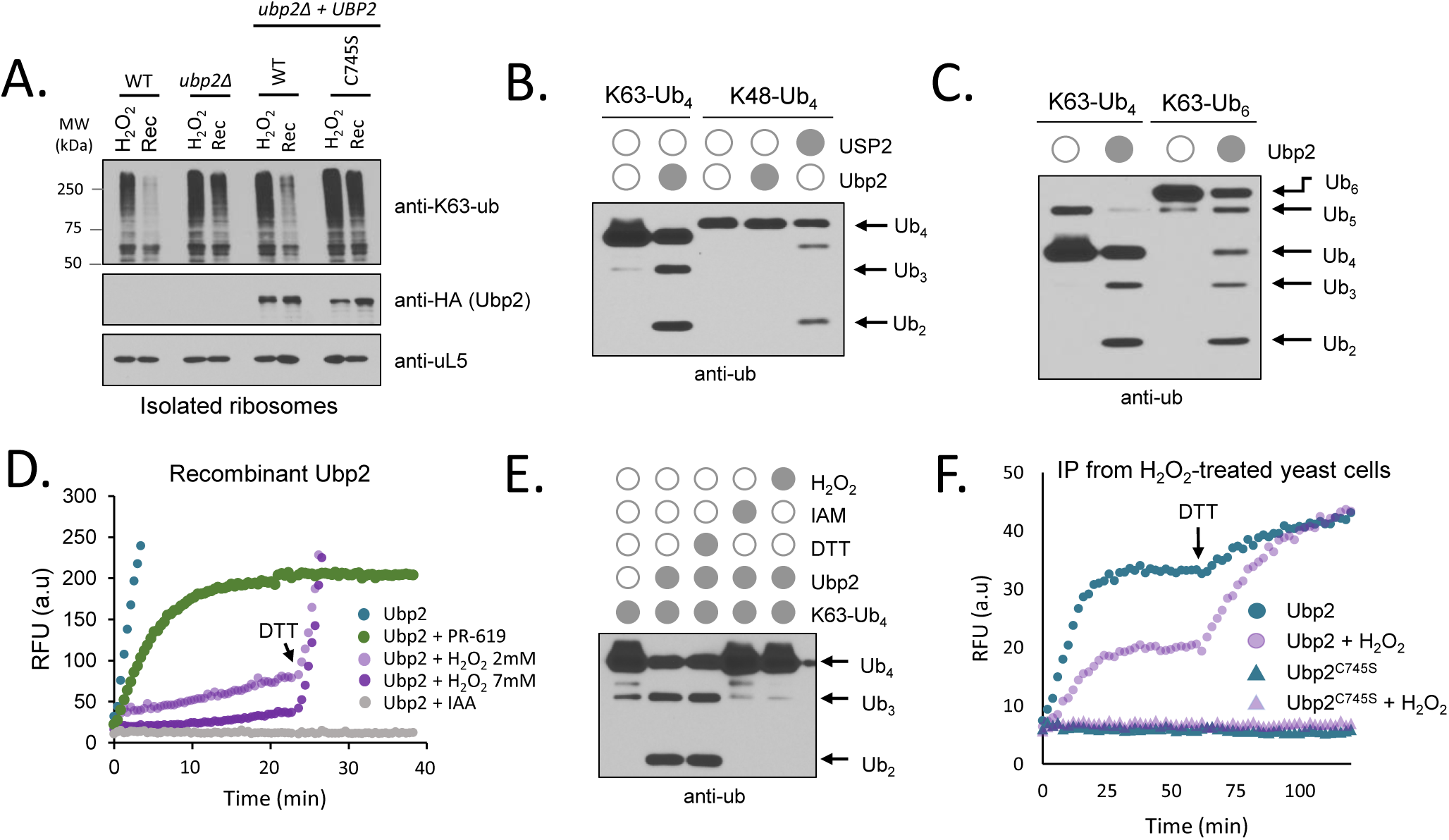
Ubp2 promotes K63-ub cleavage after stress and is reversibly inhibited by H_2_O_2_. A) Deletion and mutation to *UBP2* impair K63-ub removal from ribosomes. Immunoblot anti-K63-ub of isolated ribosomes from cells treated in the presence or absence of 0.6 mM of H_2_O_2_ for 30 min (H_2_O_2_) followed by 20 min recovery (Rec) in fresh medium. anti-uL5 was used as loading control for isolated ribosomes and anti-HA for Ubp2. B) Ubp2 preferentially cleaves K63-ub chains. Immunoblot anti-ub of synthetic tetra K63-ub or K48-ub chains incubated in the presence or absence of 3 μg of purified Ubp2 or pan DUB USP2. C) Ubp2 is a processive K63-linkage specific DUB. Immunoblot anti-ub of synthetic K63-ub chains incubated in the presence or absence of 3 μg of purified Ubp2. Arrows indicate degradation products. D) Ubp2’s activity is redox-regulated *in vitro*. Activity of recombinant Ubp2 was assessed *in vitro* using the 1.5 μM Ub-AMC fluorophore upon H_2_O_2_ treatment in the presence or absence of 20 mM IAA (cysteine alkylator) and 100 μM PR-619 (general DUB inhibitor). 10 mM of DTT (reducing agent) was added after 25 min of incubation. AMC fluorescence was recorded at 445 nm with excitation at 345 nm. E) Ubp2 deubiquitinating activity is impaired by H_2_O_2_. Immunoblot anti-ub of synthetic tetra K63-ub chains (250 ng) incubated in the presence or absence of 3 μg purified Ubp2 with the addition or not of 7 mM H_2_O_2_, 10 mM DTT, or 10 mM IAA. F) Ubp2 activity is redox-regulated in yeast cells under stress. Ubp2-HA and Ubp2^C745S^-HA were immunoprecipitated from yeast cells untreated or treated with 0.6 mM H_2_O_2_ for 30 min. Ubp2 activity was assessed *in vitro* using the 1.5 μM Ub-AMC fluorophore as in *D*. 20 mM of DTT was added after 60 min as indicated by the arrow to restore Ubp2 activity. Fluorescence was recorded at 445 nm with excitation at 345 nm. Reactions shown in *B*, *C*, and *E* were incubated for 1h at 30°C prior SDS-PAGE and immunoblotting.

### Ubp2 has different sensitivity to peroxides

To further understand the redox regulation of Ubp2, we tested whether Ubp2 activity was responsive to other peroxides beyond H_2_O_2_. To different extents, we observed that organic peroxides are also able to induce the accumulation of K63-ub chains in yeast (Fig. 2*A*). As DUBs are a diverse group of enzymes, their sensitivity to ROS can vary substantially. We first showed that cellular DUB activity was more affected by the organic peroxides cumene (CHP) and *tert-*butyl hydroperoxide (t-BHP) than H_2_O_2_ at our standard concentration of 0.6 mM, with CHP promoting the strongest effect (Fig. 2*B*, left). At 2.5 mM, organic and inorganic peroxides had a similar and acute effect in the cellular global DUB activity (Fig. 2*B*, right). These results disagreed with our previous findings that showed higher levels of K63-ub under H_2_O_2_ when compared to organic peroxides (Fig. 2*A*). Therefore, we tested whether Ubp2 itself would be more sensitive to H_2_O_2_, while cellular DUB activity would be more affected by organic peroxides. Using purified Ubp2, we confirmed that H_2_O_2_ promoted a strong reduction of Ubp2 activity (∼60%), while the organic peroxide treatments showed only a mild reduction at the same concentration (Fig. 2*C*, ∼15-30%). Upon increasing concentrations, organic peroxides could promote a stronger inhibition of Ubp2 activity (Fig. S3, *A and B*). Thus, our findings show that Ubp2 can be redox-regulated by organic and inorganic peroxides, and its sensitivity correlates to the levels of K63-ub chain observed in yeast cells (Figs. 2*A and C*, S3, *A and B*). Finally, because several DUBs are cysteine proteases (28), this redox regulation could affect several ubiquitin-mediated pathways in eukaryotic cells at once. Thus, we tested whether Cezanne, a K11-linkage specific DUB involved in cell cycle regulation (29), could also be redox-regulated by peroxides. Indeed, we also observed an increased sensitivity of the catalytic domain of Cezanne (Cezanne^CAT^) to H_2_O_2_, in comparison to t-BHP and CHP (Figs. 2*D*, S3*C*). Cezanne^CAT^ activity can also be rescued by DTT (Fig. S3). Therefore, the redox mechanisms that we are exploring for Ubp2 could have further implications for ubiquitin dynamics in a plethora of eukaryotic pathways that are responsive to ROS.

**Figure 2.**
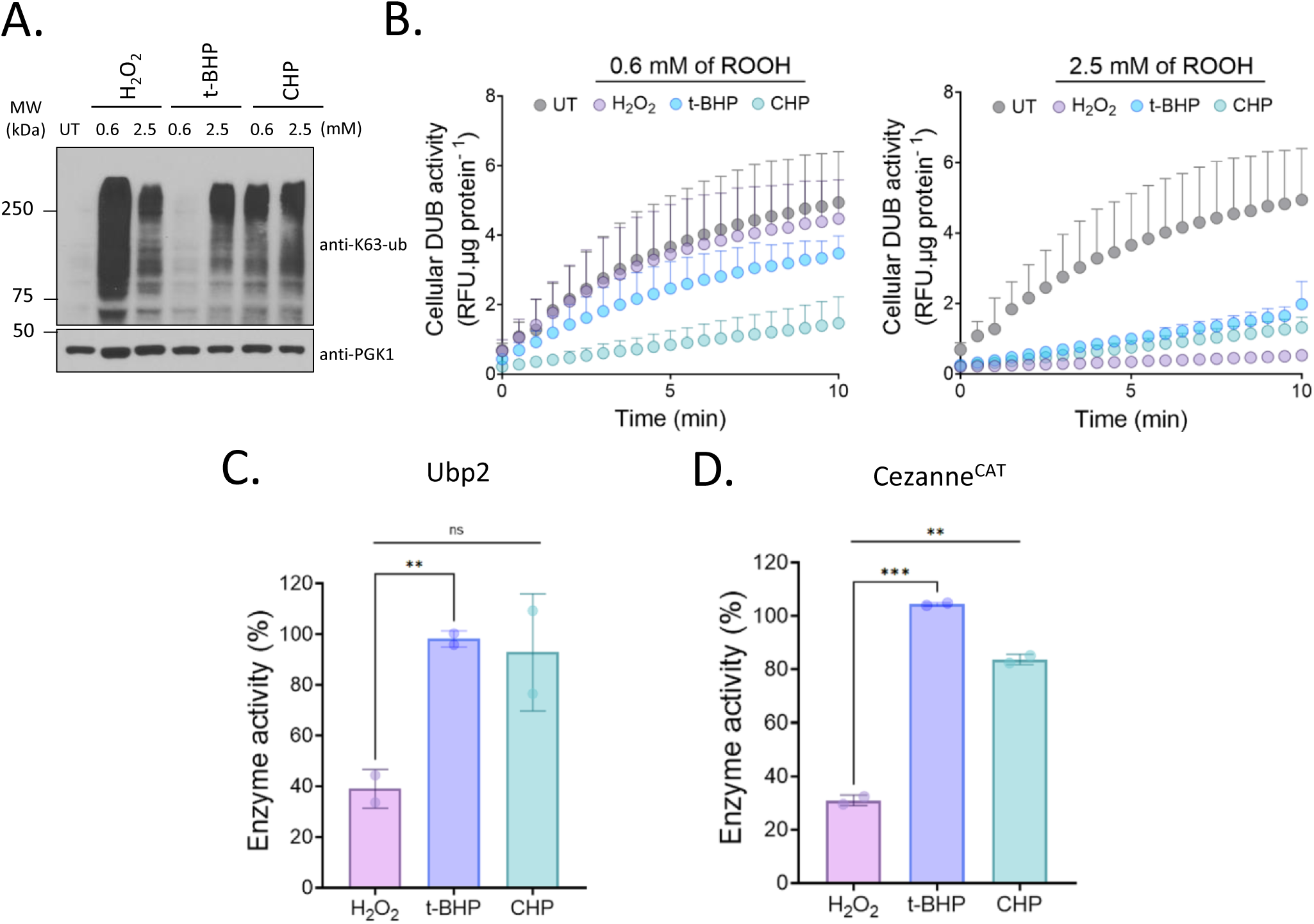
Ubp2 is particularly sensitive to H_2_O_2_. A) Accumulation of K63-ub chains is differentially induced by H_2_O_2_ and organic peroxides. Immunoblot anti-K63-ub from WT cells exposed to 0.6 mM and 2.5 mM of H_2_O_2,_ or organic peroxides *tert-*butyl (t-BHP) and cumene hydroperoxide (CHP) for 30 min. Anti-PGK1 was used as a loading control. B) Cellular deubiquitinating activity is differentially affected by H_2_O_2_ and organic peroxides. WT cells untreated (UT) or treated with either 0.6 mM (*left*) or 2.5 mM (*right*) of peroxides (ROOH) for 30 min were lysed and DUB activity determined. Activity was assessed over time using 0.75 µM of the fluorogenic substrate Ub-Rho. Rho fluorescence was recorded at 535 nm with excitation at 485 nm. C and D) Ubp2 and Cezanne^CAT^ are sensitive to H_2_O_2_. *C*, Purified Ubp2 (70 ng) and *D*, Cezanne^CAT^ (3 µg) were incubated with 500 µM H_2_O_2_, t*-*BHP, and CHP for 5 min and activity was assessed as described above. Rho fluorescence was calculated as percentage of the activity of untreated Ubp2 or Cezanne^CAT^. Bar graphs show mean values ± SD for the biological replicates. Significance was calculated using an unpaired two-tailed Student’s t-test where \**p* <0.05, ** *p* <0.005, *** *p* <0.0005, **** *p* < 0.0001 and ns considered non-significant.

### Ubp2 is reactivated during the stress recovery

Although we observed that Ubp2 activity can be reversibly regulated by ROS (Fig. 1*D and F*), it remained unclear whether this process is required to reverse the K63-ub chains that accumulate under stress. As the intracellular fate of oxidized Ubp2 is unknown, the reversal of K63-ub chains during stress recovery could be mediated either by Ubp2 reactivation or by *de novo* Ubp2 synthesis. Using the translation inhibitor cycloheximide (CHX), we first showed that the degradation of Ubp2 is not enhanced upon stress induction and Ubp2 remains present and abundant during the time required for deubiquitination (Fig. 3*A*). Moreover, yeast cells were able to cleave K63-ub chains even in the presence of CHX, suggesting that *de novo* synthesis of Ubp2 is not required for K63-ub reversal (Fig. 3*A*). Accordingly, we showed that cells treated with CHX can regain their DUB activity during the stress recovery, further indicating that redox reactivation of DUBs also occurs in the cellular context (Fig. 3*B*). Thus, our data suggest that inactivation and reactivation of Ubp2 are critical to control the levels of K63-ub chains following H_2_O_2_ stress. Although cysteine oxidation is largely a chemical process (30), intracellular reduction of these residues commonly relies on the glutathione or thioredoxin system as the main thiol-based antioxidant pathways in eukaryotes (31, 32). Therefore, we tested whether alterations of the cellular redox balance could also affect the dynamics of K63-ub during stress. Double deletions of thioredoxins (*TRX1/TRX2)*, glutaredoxins (*GRX1/GRX2) and* single deletions of thioredoxin peroxidases *(TSA1 and TSA2)* did not show sustained high levels of K63-ub chains during recovery (Figs. 3*C*, S4). However, deletion of glutathione reductase (*GLR1*) consistently impaired cell’s ability to reverse K63-ub chain during stress recovery (Fig. 3*C*). Our findings show that Ubp2 is reactivated during the recovery phase of stress in a redox-dependent manner, and suggest that cell’s reductive capacity mediated by NADPH-dependent redox systems participates in this process.

**Figure 3.**
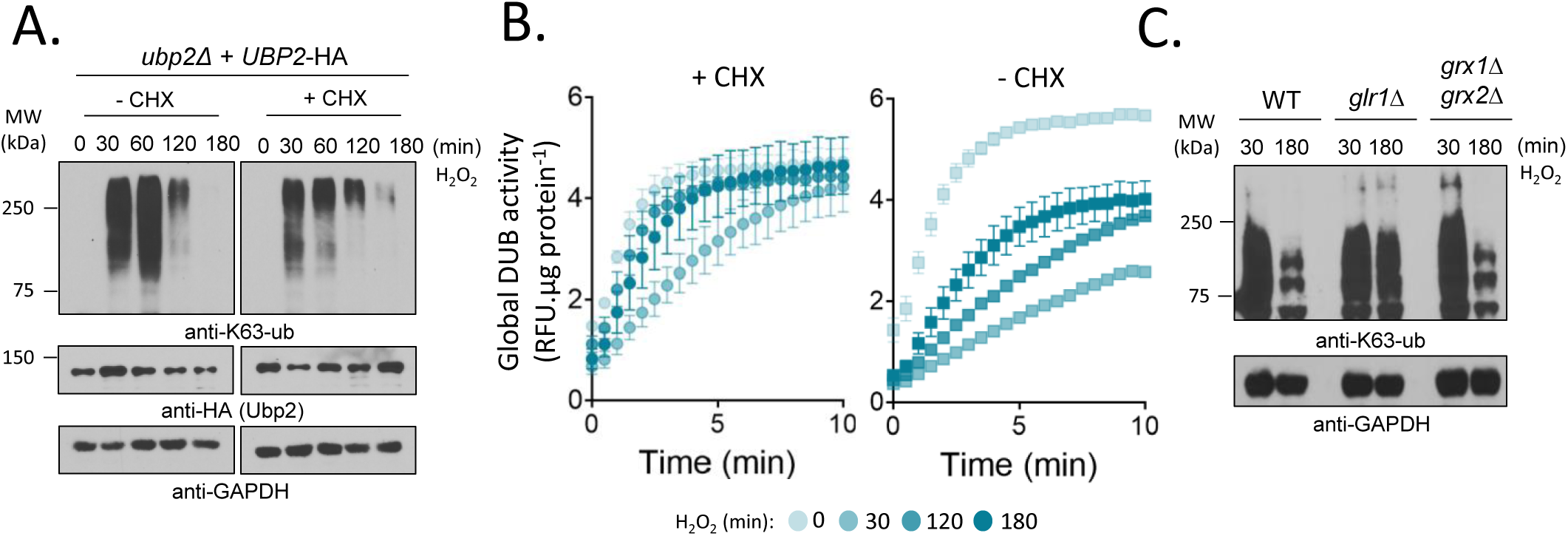
Ubp2 is reactivated during stress recovery. A) K63-ub chains are reversed independently of translation. Immunoblot anti-K63-ub from cells exposed to 0.6 mM H_2_O_2_ in the presence or absence of 150 μg/ml cycloheximide (CHX). anti-HA shows Ubp2 levels and anti-GAPDH was used as a loading control. B) Cellular deubiquitinating activity recovers from stress independently of translation. DUB activity from WT cells exposed to 0.6 mM H_2_O_2_ was assessed over time using 0.75 µM Ub-Rho in the absence (*left*) or presence (*right*) of 150 μg/ml CHX. Rho fluorescence was recorded at 535 nm with excitation at 485 nm. C) Reversal of K63-ub chains relies on cellular antioxidant systems. Immunoblot anti-K63-ub from strains incubated with 0.6 mM H_2_O_2_. anti-GAPDH was used as a loading control.

Next, we investigated the role of Ubp2 cysteine residues (Cys) on its redox regulation. Beyond their catalytic cysteine, thiol-dependent enzymes can use additional Cys to aid on their catalytic cycle and redox regulation (30, 31). Ubp2 has 13 cysteine residues, and analysis of its 3D model from AlphaFold2 (33, 34) predicted that the cysteines C821 and C944 are in proximity to the catalytic site, which could foster the formation of disulfide bonds upon conformational changes (Fig. 4*A*). By running non-reducing gels, we observed that Ubp2 forms H_2_O_2_-dependent disulfide bonds (Fig. S5*, A*-*C*) that are fully reduced by DTT (Fig. S5*C*). Mutational analysis revealed that the disulfide bond formation was abrogated in the Ubp2^C754S^ and Ubp2^C944S^ mutants following H_2_O_2_ treatment, indicating that these residues are involved in disulfide bond formation (Fig. 4*B*). To determine the effect of these cysteine residues in Ubp2 activity, we evaluated cellular growth of strains carrying selective Ubp2 mutations through cellular sensitivity to the proteotoxic agent L-Azetidine-2-carboxylic acid (ADCB). Functional Ubp2 provides resistance to ADCB by deubiquitinating and recycling selective ADCB transporters (13, 35). First, we showed that cells expressing the catalytic dead Ubp2^C745S^ mutant are highly sensitive, while cells carrying WT Ubp2 show tolerance to ADCB (Fig. 4*C*). Next, we tested whether mutation to other cysteine residues affects cell’s sensitivity to ADCB. While we observed a mild effect for the double mutation C821S/ C944S to serine, mutations to cysteine C621 and C645 show no change in ADCB sensitivity (Fig. 4*C*). These results suggest that the presence of C821 and C944 is important to support Ubp2 structure or function. However, it was still unclear whether these residues participate in K63-ub reversal as part of the RTU. Our results show that the strain expressing Ubp2 C821S/C944S were still able to remove K63-ub chains in a timely fashion, at least under the conditions tested (Fig. 4*D*). Thus, our data shows that Ubp2 is reactivated upon stress cessation and is able to form disulfides bonds in the presence of ROS, which correlates to the levels of K63-ub chains that accumulate under stress.

**Figure 4.**
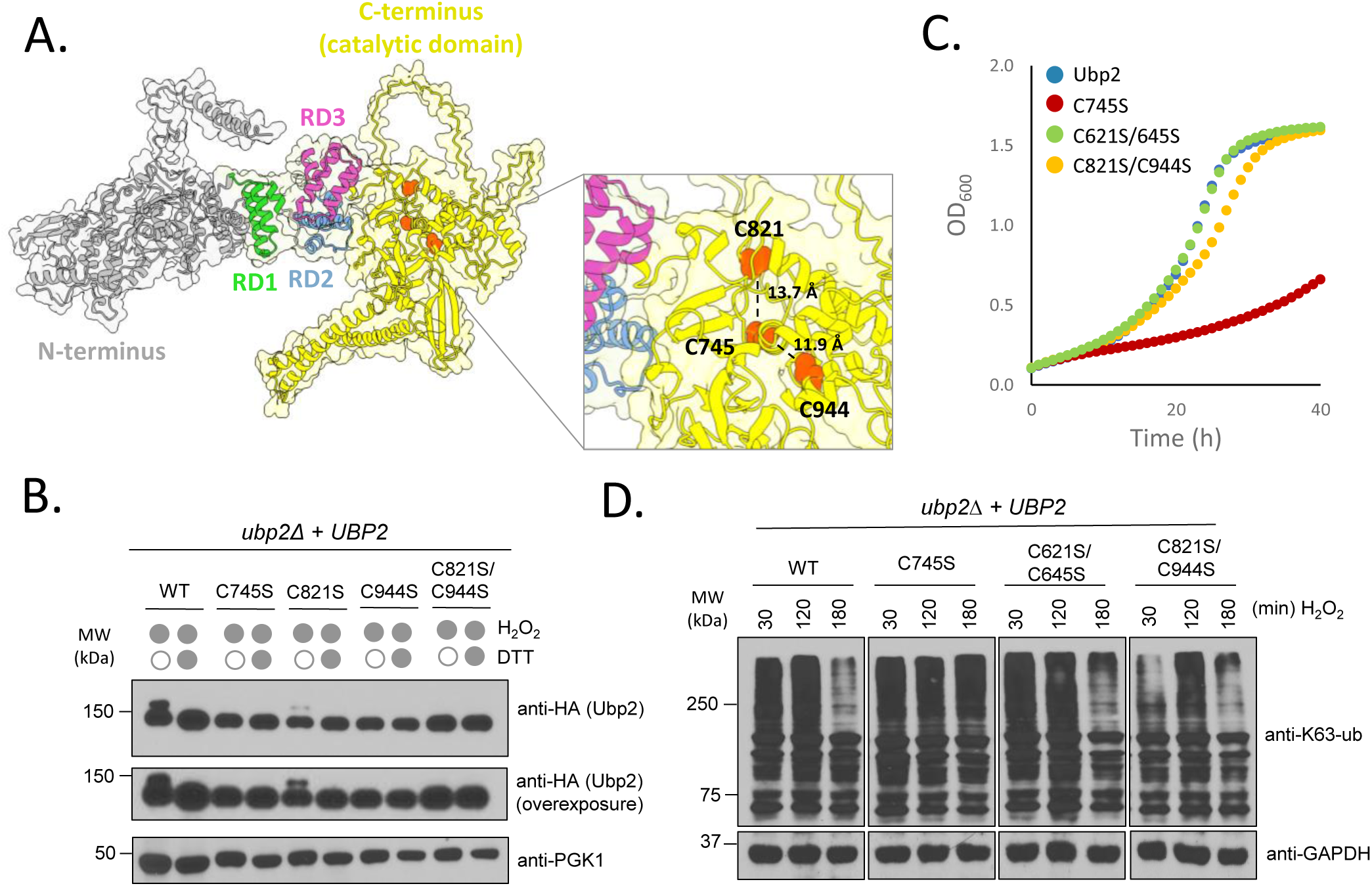
Ubp2 catalytic cysteine forms disulfide bonds under stress. A) AlphaFold2 structural 3D model of Ubp2 (ID: Q01476). N-terminus (grey), Repeated domain (RD) 1 (green), RD2 (blue), RD3 (pink) and C-terminus (yellow). Inset highlights in orange displays the catalytic cysteine (C745) and neighboring cysteine residues (C821 and C944) in the catalytic domain with its predicted molecular distances. B) Mutation to catalytic cysteine C745 and C944 inhibits formation of disulfide bond under stress. Immunoblot anti-HA shows Ubp2 levels from cells treated with 0.6 mM H_2_O_2_ for 30 min. Lysates were incubated in the presence or absence of 20 mM DTT prior to immunoblotting. anti-PGK1 was used as a loading control. C) Cellular sensitivity to the proteotoxic agent ADCB. Cell growth was monitored through absorbance (OD_600_) over time upon addition of 100 µg/ml ADCB. Ubp2^FL^ and Ubp2^C745S^ were used as positive and negative control, respectively. D) Comparative reversal of K63-ub chains during stress recovery. Immunoblot anti-K63-ub from cells expressing Ubp2-HA or its cysteine mutants upon treatment with 0.6 mM H_2_O_2_ for the designated times. anti-GAPDH was used as a loading control.

### Ubp2 activity is regulated by a series of repeated domains

We have identified key cysteine residues involved in the redox regulation of Ubp2, however additional functional elements are necessary to drive its deubiquitinating function. Ubp2 is comprised of a non-conserved N-terminus extension, three repeated domains (RDs) and a conserved C-terminus UBP/USP catalytic domain (Figs. 4*A*, 5*A*). While it is known that Ubp2 plays a role in several pathways mediated by K63-ub (5, 36), the protein domains and residues that are critical for its activity have only been broadly characterized in the context of its interaction with the E3 Rsp5 and the co-factor Rup1 in vesicle trafficking (37). As reversal of K63-ub that accumulate under stress works independently of Rup1 (Fig. S6*A*), we sought to characterize the role of Ubp2’s domains in the RTU. To determine the functional role of Ubp2 domains, we turned again to our ADCB experiment that allow us to test several experimental conditions in parallel. We observed that expression of the N-terminus only (Ubp2^Nterm^) or the catalytic domain only construct (Ubp2^CAT^) led to cellular sensitivity towards ADCB (Fig. 5*B*). Ubp2^CAT^ still retained partial activity against the fluorogenic substrate Ub-AMC (Fig. S6*B*), suggesting that additional domains are important for Ubp2 cellular function. By producing additional truncated versions, we showed that cells expressing Ubp2 lacking its N-term but expressing its three RDs and C-terminus (Ubp2^RD1-3^) are sensitive to ADCB (Fig. 5*C*). Surprisingly, upon deletion of RD1 (Ubp2^RD2-3^), cells resistance to ADCB was rescued closer to WT levels (Fig. 5*C*). We then removed Ubp2 RD2 (Ubp2^RD3^), which also showed increased sensitivity to ADCB (Fig. 5*C*), suggesting that RD2 and the catalytic domain are required to promote resistance against ADCB. To test whether RD2 alone would be sufficient to support Ubp2 function, we created new constructs and showed that cells expressing RD2 with the catalytic domain (Ubp2^RD2^) remain sensitive to ADCB (Fig. 5*D*). Thus, our results indicate that RD2 is required but not sufficient to regulate Ubp2 activity. Addition of RD1 (Ubp2^RD1-2^) makes cells more sensitive to ADCB again (Fig. 5*D*). The presence of RD1 consistently reduces Ubp2 activity, suggesting that it might serve as an inhibitory domain that could reduce the exposure of key catalytic residues to the substrate.

**Figure 5.**
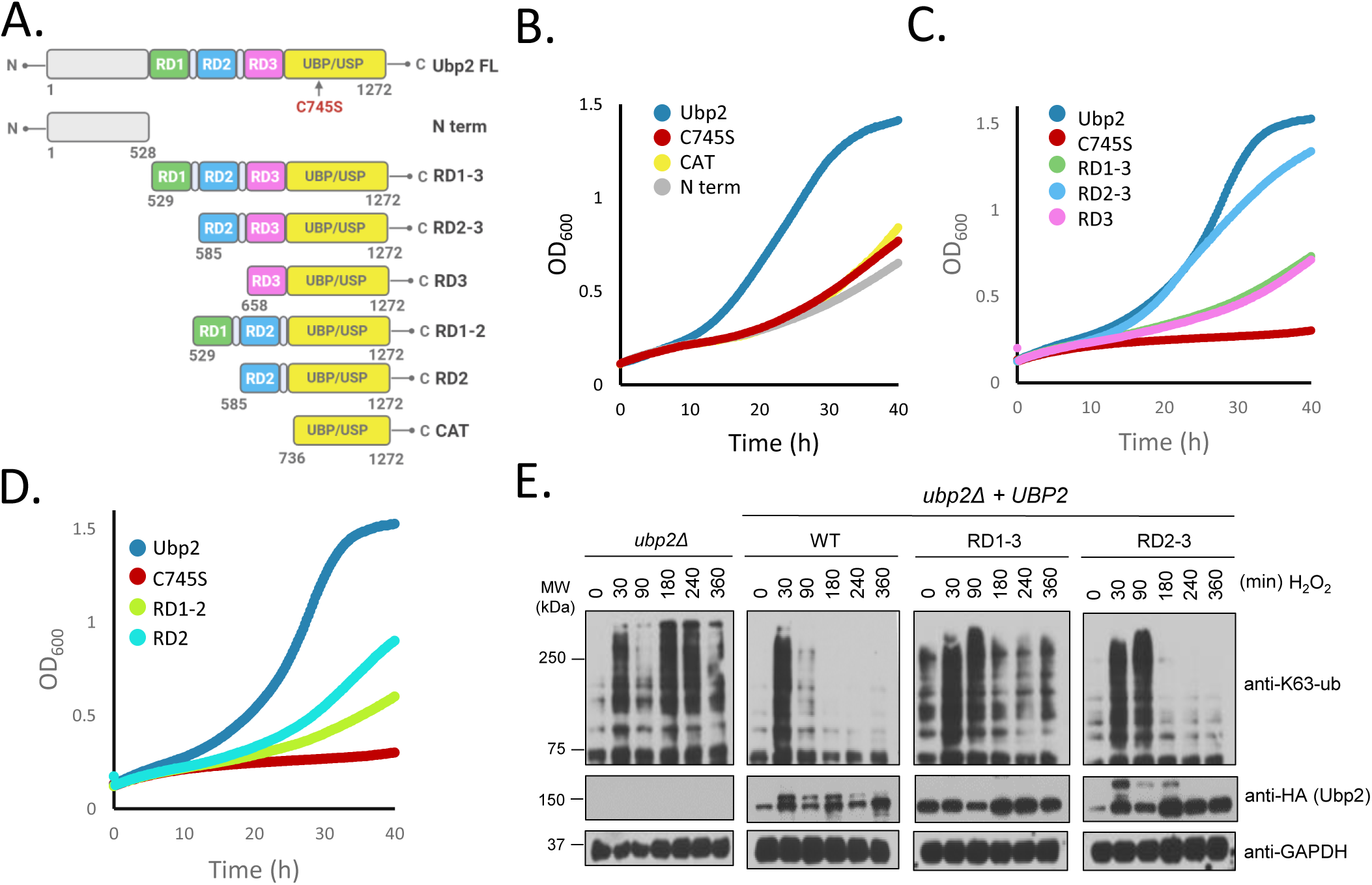
Ubp2 is regulated by a series of repeated domains. A) Schematic of Ubp2 domain organization and truncations created in this study. Ubp2 is comprised of a non-conserved N-terminus, three repeated domains (RDs), and a conserved C-terminus UBP/USP catalytic domain. Created with BioRender.com B-D) Ubp2’s RDs modulate cell’s resistance against proteotoxic agent ADCB. ADCB sensitivity growth curves for the Ubp2 truncations as labeled above. Cell growth was monitored through absorbance (OD_600_) over time upon addition of 100 µg/ml ADCB. Ubp2^FL^ and Ubp2^C745S^ were used as positive and negative control in all growth curves, respectively. E) Ubp2’s RDs regulate K63-ub chain removal during stress recovery. Immunoblots anti-K63-ub of *ubp2Δ* cells expressing WT Ubp2, Ubp2^RD1-3^, and Ubp2^RD2-3^ in the presence or absence of 0.6 mM H_2_O_2_. anti-GAPDH was used as loading control.

Following our initial functional screen of Ubp2’s RDs, we asked whether these regulatory domains also affected the reversal of K63-ub chains that accumulate in response to stress. We have not identified any relationship of these domains with Ubp2’s capacity to bind or associate with ribosomes (Fig. S6, *C and D*), but they indeed affected the ability of cells to reverse K63-ub chains. In agreement with the ADCB experiments, cells expressing Ubp2 containing all repeated domains (Ubp2^RD1-3^) were inefficient in reversing K63-ub chains, while deletion of RD1 (Ubp2^RD2-3^) also rescued the phenotype and contributed to reversal of these ubiquitin chains (Fig. 5*E*). Supporting the notion that RD1 might play an inhibitory effect, we observed that Ubp2^RD2-3^ forms high levels of disulfides after H_2_O_2_ (Figs. 5*E*, S5*A*), which are also prevented by the presence of RD1. Here we showed that Ubp2 activity is regulated by a series of repeated domains that can control cell resistance to ADCB and the levels of K63-ub chains in the RTU.

### Ubp2 supports the re-establishment of translation upon stress cessation

We have previously determined that K63-ub that accumulates in response to oxidative stress modifies ribosomes and pauses translation at the elongation stage (5, 7, 10). Following oxidative stress induction, Ubp2 is required for ubiquitin reversal, however, it remains unclear how Ubp2 regulates translation through its deubiquitinating activity. To start to elucidate the role of Ubp2 in translation, we first showed that purified Ubp2 is able to deubiquitinate isolated ribosomes *in vitro and* the deubiquitination of ribosomes by Ubp2 can also be impaired by H_2_O_2 and_ reversed by DTT (Fig. 6*A*). Using polysome profiling, we showed that Ubp2 remains bound to the ribosomes in the presence or absence of stress and cells lacking Ubp2 retained high levels of K63-ub chains in the monosome and polysome fractions during stress recovery (Fig. 6, *B and C*), further suggesting that Ubp2 is responsible for removing K63-ub chains from elongating ribosomes. Because of the role of K63-ub in ribosome pausing (7, 10), we hypothesized that cells lacking Ubp2, would show impaired resumption of protein synthesis during stress recovery. Using an inducible GFP-based reporter, we showed that both strains, the WT and *ubp2Δ,* have GFP expression inhibited by H_2_O_2_ (Fig. 6*D*). However, while WT cells restore GFP synthesis after ∼3h following stress induction, GFP production remains significantly lower in *ubp2Δ* (Fig. 6*D*). Our findings suggest that Ubp2 plays a role in translation resumption following stress, however it was unclear whether the results observed for this GFP reporter could be extrapolated at the proteome level.

**Figure 6.**
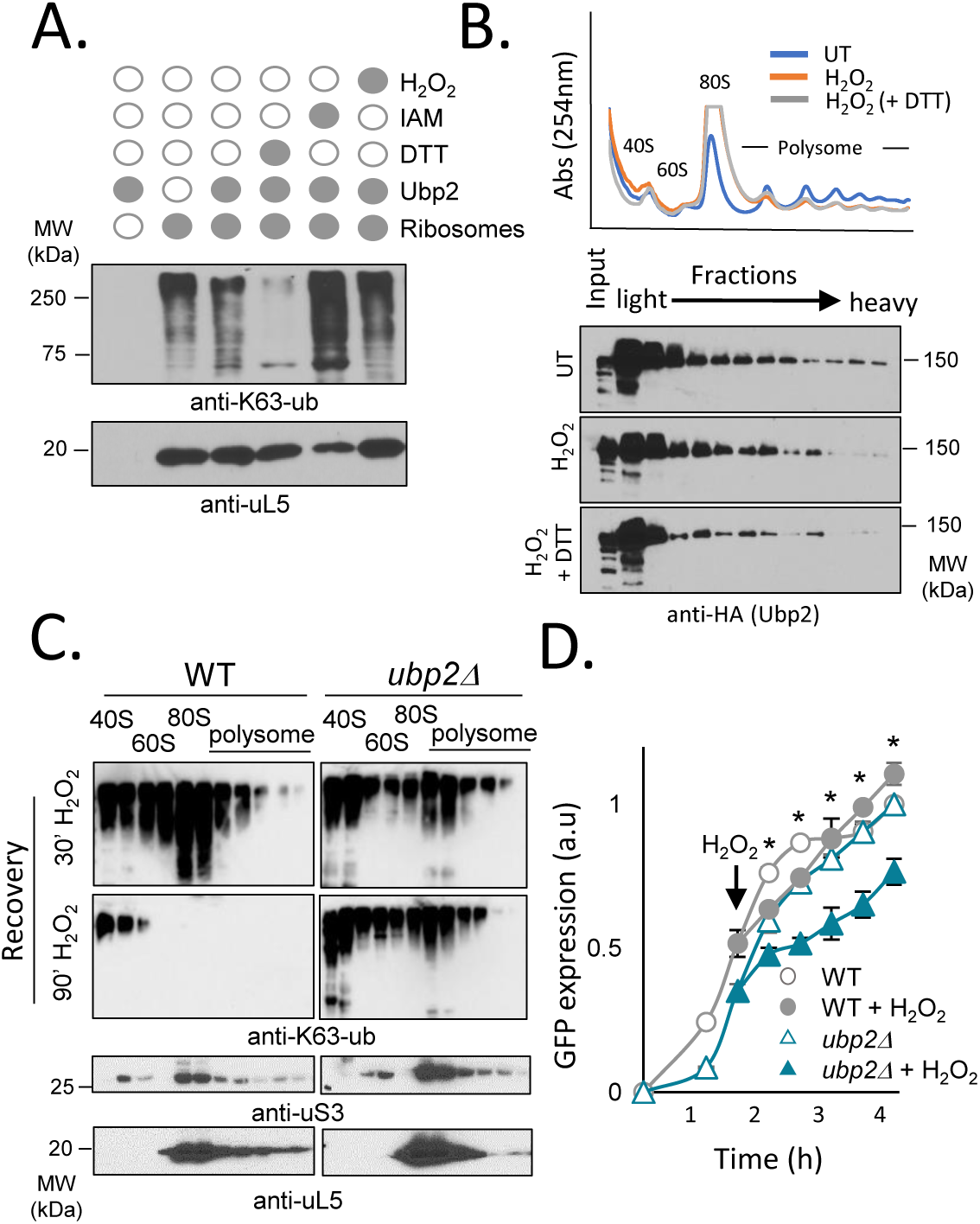
Ubp2 deubiquitinates ribosomes and regulates protein synthesis. A) Ubp2 deubiquitinating activity against ribosomes is redox-regulated *in vitro*. Immunoblot anti-K63-ub of ribosomes (40 µg) isolated from *ubp2Δ* cells. Ribosomes were incubated in the presence or absence of 10 mM H_2_O_2_, 10 mM IAA (cysteine alkylator), 10 mM DTT (reducing agent), and purified recombinant Ubp2 (5 µg) for 1h at 30°C at 300 rpm. anti-uL5 was used as a loading control. B) Ubp2 is associated with ribosomes in the presence or absence of H_2_O_2_. Immunoblot anti-HA to detect Ubp2 levels from polysome profiling fractions from WT cells untreated or treated with H_2_O_2_ for 30 min. C) *ubp2Δ* cells present delayed K63-ub reversal from ribosomes during stress recovery. Immunoblot anti-K63-ub of ribosomes fractions isolated from WT and *ubp2Δ* upon treatment with 0.6 mM H_2_O_2_ for 30-(*top*) and 90 min (*bottom*), n =1. anti-uS3 and anti-uL5 were used as a loading control for 40S and 60S ribosome subunit, respectively. D) Fluorescence of GFP-reporter in Ubp2-HA and *ubp2Δ* cells was analyzed as a proxy for translation activity and translation recovery. GFP expression was induced in –Met medium followed by the addition of 0.6 mM H_2_O_2_ at 100 min as shown by the arrow. Significance was calculated using an unpaired Student’s t-test between WT/WT H_2_O_2 and_ *ubp2Δ*/*ubp2Δ* H_2_O_2_ where comparisons with *p*> 0.05 were considered non-significant (ns) and samples with *p* <0.05 were considered significant (*).

To further investigate Ubp2’s role in restoring protein synthesis, we performed quantitative proteomics to understand how Ubp2 contributes to a global mechanism of translation regulation during different phases of the oxidative stress response. For this purpose, we cultivated WT and *ubp2Δ* cells and measured protein abundance in cells untreated or incubated with H_2_O_2_ for 30- and 120-min using label-free mass spectrometry. We chose 120 min after stress induction because at this time, cells start to reverse the K63-ub chains that accumulated under stress and resume translation that had been inhibited following H_2_O_2_ treatment (Figs. 3*A*, and 6*D*). Our processed dataset is comprised of 4422 proteins representing ∼75% the yeast proteome (38, 39). For a small number of proteins (179 proteins, 4.0% of the dataset) that presented three or fewer missing values across the 18 samples (*i.e.*, 2 strains, 3 time points, 3 replicates), intensity values were imputed using k-nearest neighbor method (40). To confirm that cells were responding to H_2_O_2_, we showed that both strains significantly enhanced the expression of key antioxidant enzymes such as thioredoxin Tsa2 and catalase Ctt1 after stress induction (Fig. S7*A*). When accounting for basal differences between strains, we observed that after 30 min of stress, 23 proteins are significantly down-regulated (<1.5 fold) in the *ubp2Δ* compared to the WT (Fig. S7*B*). Gene ontology (GO) analysis revealed that proteins involved in mitotic spindle and membrane proteins are significantly down-regulated in the *ubp2Δ* background (Fig. S7*D*). When focusing on the recovery phase (30 to 120 min), and accounting for basal differences among strains, we observed that 8 and 27 proteins are significantly down- or up-regulated, respectively, in the *ubp2Δ* strain (Fig. S7*C*). GO analysis confirmed that molecular functions related to exocytosis, vacuolar membrane, and mitosis remain up even 120 min after H_2_O_2_ addition to media (Fig. S7*E*).

Our results using GFP as a reporter protein indicated that Ubp2 is important for the re-establishment of protein synthesis during the recovery phase of stress (Fig. 6*D*). When comparing 120 min to our initial time point, we observed that 341 proteins are still differentially expressed in *ubp2Δ* in contrast to 186 in the WT (Fig. S7, *F and G*). Of that, 90 proteins were shared among strains (Fig. S7*G*). Although only a fraction of the proteome is substantially different, our results suggest that deletion of *UBP2* affects specific cellular pathways, however other meaningful physiological changes might be present below our fold-change cutoff. Our findings related to mitosis and mitotic spindle, motivated us to inspect the distribution of functional protein groups that are directly associated with cellular growth/division and stress response. We first observed that proteins from the GCN4 regulon, which are expressed as part of the integrated stress response (41), are significantly lower in *ubp2*Δ (Fig. 7*B*). This suggests that translation dysregulation might also affect cells capacity to amount a proper antioxidant defense. We also observed that ribosomal proteins are largely enriched among the up-regulated proteins in WT cells (Fig. 7*A*). However, these proteins are mostly represented amongst the down-regulated proteins and significantly lower in *ubp2*Δ (Fig. 7*B and C*). As expression of these proteins is controlled by the TORC1 pathway (42), their levels are highly correlated with cellular growth and fitness (43). Interestingly, we observed that only 16.7% and 9.1% of the proteins that are up- or down-regulated, respectively, overlap comparing WT and *ubp2Δ* strains, which suggests that these cells are mounting different cellular responses to restore proteostasis (Fig. 7*D*). These findings support our hypothesis that Ubp2 plays an important role is rescuing proteostasis and cellular growth following stress. Taken together, our results support a model where Ubp2 is an integral part of the RTU, in which its redox regulation determines the dynamics of ribosome ubiquitination that is required to achieve a proper stress recovery.

**Figure 7.**
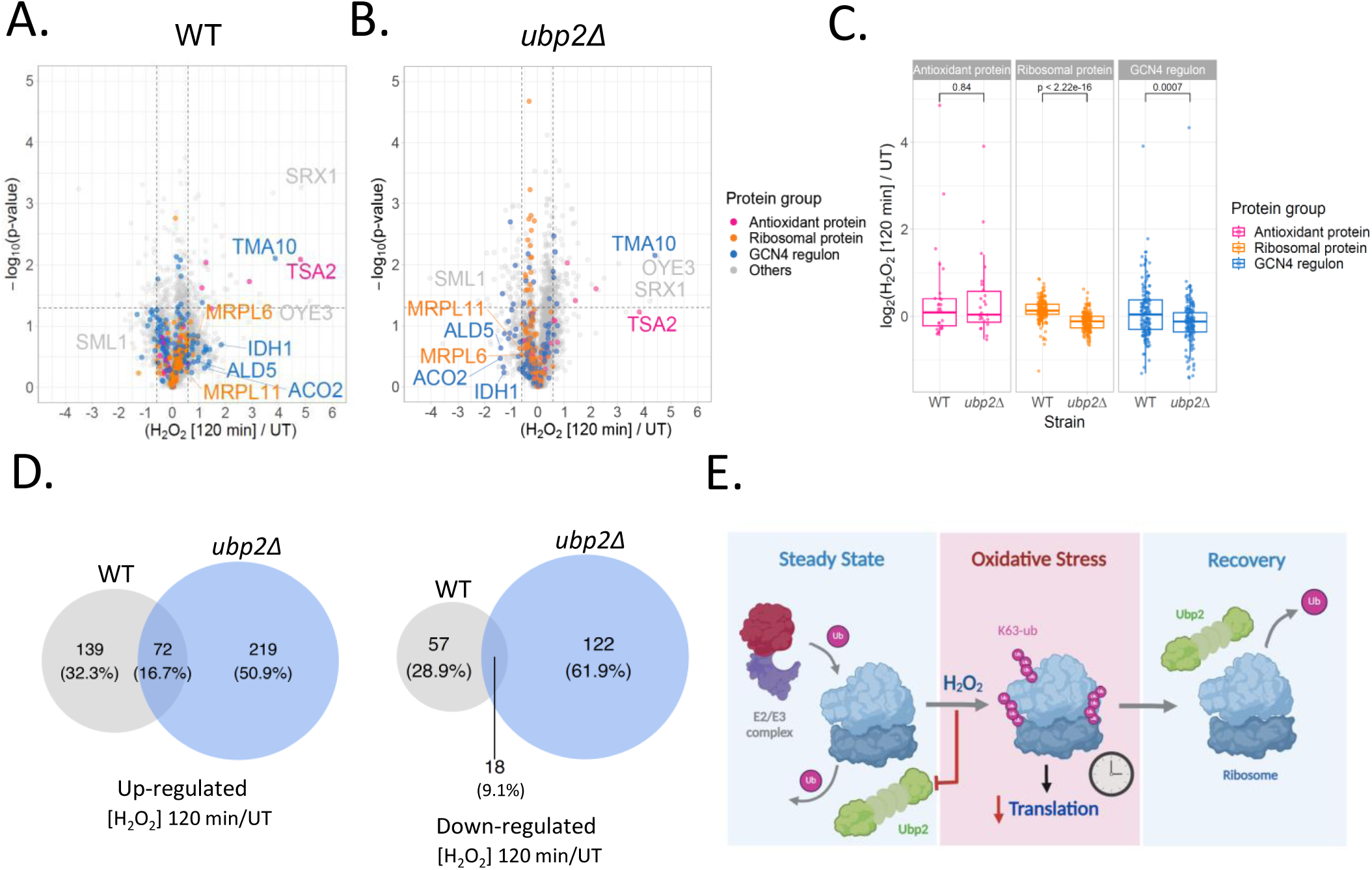
Ubp2 supports translation reprogramming following stress. A-B) Volcano plots displaying changes in protein levels for WT and *ubp2Δ* strains comparing 120 min after H_2_O_2_ treatment to untreated (UT) condition. Proteins are color-coded based on their subgroups: antioxidant proteins (pink), ribosomal proteins (orange), GCN4 regulon (blue) and others (grey). The horizontal dashed line indicates significance (p < 0.05), while the vertical dashed lines represent a fold change of ±1.5. C) Box plots quantification for proteins belonging to the three functional groups as above. p-values derived from unpaired Student’s t-test. D) Venn diagrams showing the proteins up- (*left*) and down-regulated (*right*) in WT and *ubp2Δ* cells between 120 min after H_2_O_2_ treatment and untreated (UT) conditions. E) Schematic model for Ubp2 role in the RTU during steady state, stress condition, and stress recovery. Created with BioRender.com

## Discussion

Here we showed that redox regulation of the deubiquitinating enzyme Ubp2 is critical for the modulation of translation under stress. Our results demonstrated that Ubp2 is able to cleave K63- ub chains *in vitro and* in the cellular context (Fig. 1*, A-C*) and its activity can be reversibly regulated by peroxides (Fig. 1*D*). Although details on the molecular regulation of Ubp2 remain to be further elucidated, our work revealed the importance of several Ubp2’s domains and amino acid residues on its redox regulation and activity (Figs. 4 and 5). This work opened several directions of research as a large fraction of eukaryotic DUBs are cysteine proteases (20, 23, 24) and have the potential to be redox-regulated under stress. Our findings using different peroxides (Fig. 2*B*) further demonstrate the nuances of the oxidative stress response, suggesting that different DUBs might be preferentially regulated under specific ROS conditions, triggering selective pathways. In this pathway, the redox balance of the cell and key thiol specific enzymes (Figs. 3*C*, S4), likely function in a cascade that donates electrons for Ubp2 reduction and reactivation.

The indications that Ubp2 repeated domains could have inhibitory or activation functions (Fig. 5, *C-E*) adds to this complexity and further research must be conducted to understand the structural and functional conservation of these repeated domains across the evolutionary scale. While Ubp2 RD1 show an inhibitory effect, RD2 seems required but not sufficient to drive Ubp2 function. In this context and based on AlphaFold2 structural predictions, RD3 might possess a structural role in positioning RD2 towards the catalytic residues (Fig. 4*A*). We also showed that the catalytic cysteine C745 is able to form a disulfide bond with C944 under stress (Fig. *4*B) and mutations affect Ubp2 activity (Fig. 4*C*). However, the role of this disulfide in the RTU remains to be elucidated. Although cells expressing Ubp2^C944S/C821S^ are able to reverse K63-ub chains, it is unclear under different experimental conditions what fraction of Ubp2 remains active, what fraction forms disulfides, and what fraction can be hyperoxidized, which would prevent its redox recycling. These findings support our conclusions that the activity of Ubp2 rely on these series of repeated domains and future experiments should address the structural details by which Ubp2 recognizes and binds to ribosomes and K63-ub chains.

One of the first steps of the RTU is the accumulation of K63-ub chains on ribosomes during stress (5, 7). We have previously shown that the E2 conjugase Rad6 and the E3 ligase Bre1 are mainly responsible for the burst of the K63-ub chains upon H_2_O_2_ exposure (11). Moreover, we have shown that Rad6 itself can be redox-regulated and form an intermolecular disulfide bond with the E1 Uba1 (11). This regulation is part of a feedback loop that controls the total amount of ubiquitin moieties added to ribosomes during the reprogramming of translation (11). In addition to the regulation of Rad6, we showed that Ubp2 plays a central role in the accumulation and reversal of these K63-ub chains (Fig. 1*A*). Our findings suggest a cycle of ubiquitination and deubiquitination in which Rad6 and Bre1 are constantly counteracted by Ubp2, which determines the amount of K63-ub chains in the system (Fig. 7*E*). Supporting this notion, we have observed that deletion of *UBP2* leads to higher levels of K63-ub chains even in the absence of stress (5). In addition, this early accumulation of K63-ub chains further indicate that Ubp2 is the first protein in the RTU to be inhibited upon H_2_O_2_, which leads to a net gain of K63-ub chains. Next Rad6 becomes inhibited, which defines the total amount of K63-ub added to ribosomes (5). Following stress cessation, Ubp2 is reactivated and is able to cleave these chains from its targets, returning translation to steady state (Fig. 7*E*). Considering that several DUBs can be redox-regulated in a similar fashion, there is the potential that many pathways beyond the RTU can be controlled by a dynamic cycle of ubiquitination and deubiquitination. This cycle can then be temporarily interrupted depending on the nature of ROS, the intensity, and the duration of the stresses to which cells are exposed.

Although ubiquitination has been traditionally related to protein degradation (44), several new proteasome-independent pathways have been uncovered (45, 46). For example, different steps in gene expression, translation, and protein quality control are known to use ubiquitin in a regulatory manner (47, 48). The regulation of transcription by ubiquitination has been heavily investigated (49, 50), but the rules of translation regulation are far less understood particularly in response to dynamic environments. As several aspects of translation control can be regulated by ubiquitin (48), DUBs are known to play crucial roles in antagonizing these pathways through the modulation of cycles of ribosomal ubiquitination and deubiquitination. In the RTU, ubiquitination and the activity of Rad6 are required to pause translation and reprogram protein production under stress (10). The ribosomes-associated quality control pathway (RQC) is also controlled by the ubiquitination of ribosomal proteins and has been suggested to rely on the DUBs OTUD3 and USP21 (51). Moreover, deubiquitination of the 40S protein eS7 by the DUB Ubp3 regulates translation efficiency and the DUB Otu2 promotes 40S dissociation from mRNA, participating in the recycling stage(52). In addition, the DUBs USP10 has been found to rescue ubiquitinated and stalled ribosomes from lysosomal degradation (53), while USP36 is required for maturation of the 40S ribosomal subunit (54). However, how these additional DUBs are regulated, how the ubiquitin dynamics in the ribosome is affected under stress, and level of cross-talk among these different pathways remains largely unknown (55). One of the key questions that remain open in the RTU is how ubiquitin pauses ribosomes under stress and how their deubiquitination allows translation to resume. Because of a preferential pause of ribosomes at the pre-translocation stage of translation elongation with sequence-specific features (7, 10), we have previously proposed that ubiquitin affects the dynamics of binding and recruitment of translation factors to ribosomes (7). However, further research will be necessary to define the molecular and structural characteristics of this pathway. Regardless, the reversible regulation of Ubp2 allows us to propose a mechanism by which ubiquitin temporarily pauses ribosomes, which are allowed to continue to translate once they are deubiquitinated (Figs.1*A*, 6*A and C*).

Throughout evolution, eukaryotic genomes have expanded and humans currently encode ∼100 DUBs with new functions, specialization, and pathway redundancy (56). Although Ubp2 is not highly conserved in humans, this surveillance mechanism mediated by Ubp2 in association with ribosomes will also support the discovery of new pathways that share similar molecular rules. Other groups have shown that a number of human DUBs can be activated or enhanced by reducing conditions (21, 27, 57), highlighting that we are just in the infancy of understanding how DUBs control cellular physiology globally in response to stress. As several diseases are caused by mutations and impairment of DUB activity (58), elucidating new mechanisms of DUB regulation can uncover new targets and support the development of new modes of therapy.

## Experimental procedures

### Yeast strains, plasmids, culture, and protein extraction

All *Saccharomyces cerevisiae* strains and plasmids used in this study are described in Supporting Table 1. The resumed proteomics analyses and metadata is in Table 2. Unless specified, yeast cells were cultivated into synthetic dextrose minimal medium (SD: 0.67% yeast nitrogen base, 2% dextrose and required amino acids) at 30°C at 200 rpm agitation. Cells were harvested at exponential phase OD_600_ 0.3-0.5. Protein extraction, ribosome isolation, polysome profile, GFP expression, and preparation for immunoblotting assays were performed as described previously (11).

### Protein expression and purification

*E. coli* BL21 and BL21-CodonPlus (DE3)-RIL were transformed with pCS (Cold-Shock induced bacterial vector)/*UBP2*-HA-TEV-His and pGEX/Cezanne^CAT^ (catalytic domain)-TEV-GST, respectively. Bacterial cells were grown until OD_600_ reached 0.6 – 0.8. Ubp2 expression was induced overnight at 16°C in the presence of 1 mM isopropyl ß-D-1-thiogalactopyranoside (IPTG) and Cezanne^CAT^ was expressed with 0.6 mM IPTG for 4h at 37°C. Ubp2’s cell lysis was carried out by incubation with 1mg/ml lysozyme for 1h at 4°C in buffer containing 50 ml Tris-HCl pH 7.5, 500 mM NaCl, 1 mM of DTT, and protease inhibitors (1 mM PMSF and 10 µM leupeptin) during 4x rounds of 2 min of sonication on ice followed by 1 min of rest. *E. coli* cells expressing Cezanne^CAT^ were lysed by sonication in NETN buffer (50 mM Tris-HCl pH 7.5, 150 mM NaCl, 0.5% IGEPAL) with 1mM DTT, 1 mM PMSF, 1 mM leupeptin, and 1 mg/ml lysozyme. Extract was cleared by centrifugation at 12,000 rpm for 30 min prior to 2h incubation with nickel affinity beads for Ubp2 purification (GOLDBIO catalog H-355) or glutathione-containing beads (GOLDBIO G-250-5) for 2h at 4°C for Cezanne^CAT^. The elution of Ubp2 was carried out with 250 mM imidazole followed by size exclusion chromatography column (SEC 200 – Cytica HiLoad 26/600 Superdex 200 pg, cat. # 28989336) in buffer containing 50 mM Tris-HCl pH 7.5, 150 mM and 1 mM DTT. Cezanne^CAT^ was eluted by cleavage of the GST tag with 15 units/ml TEV protease (SigmaT4455-1KU) in PBS buffer overnight at 4°C and desalted using a PD-10 column (GE Healthcare, cat. # 17085101) in 50 mM Tris-HCl pH 7.5, 100 mM NaCl buffer. Fractions were combined and concentrated using an Amicon centrifugal filter with 100 kDa or 30 kDa cutoff (Sigma), for Ubp2 and Cezanne^CAT^, respectively.

### In vitro ribosome deubiquitinating assay

The *in vitro* deubiquitinating assay was performed in the presence of 5 μg Ubp2, 10 mM DTT, and 40 μg of isolated ribosomes. All components were pre-incubated in reaction buffer (50 mM Tris pH 7.5, 100 mM NaCl, and 10 mM MgCl_2_) for 10 min at room temperature, before the addition of ribosomes. When specified, 7 mM H_2_O_2 and_ /or 10 mM IAA was added to the reaction prior to the addition of ribosomes. The reaction was incubated for 1h at 30°C at 300 rpm, stopped by the addition of 4X Laemmli sample buffer, and subjected to SDS-PAGE gel prior to immunoblotting.

### K63-ub chain cleavage assay

The cleavage of K63-ub chains by Ubp2 was assessed through an *in vitro* reaction containing 3 μg Ubp2, 10 mM DTT, 150 ng of tetra (LifeSensors) or 250 ng hexa K63-ub chains (R&D Systems), and reaction buffer (50mM Tris pH 7.5, and 100 mM NaCl). The reaction was incubated for 1h at 30°C at 300 rpm, stopped by the addition of 4X Laemmli Sample Buffer, and subjected to SDS-PAGE followed by immunoblotting.

### Immunoprecipitation assay

Yeast cells expressing Ubp2-HA and Ubp2^C745S^-HA were grown until OD_600_ 0.3-0.4 and challenged with 0.6 mM H_2_O_2_. Lysis was carried out in 50 mM Tris-HCl, pH 7.5, 100 mM NaCl, 5 mM MgCl_2_, 20 mM KCl as described before (11). 25 µl of the normalized lysate was added to Pierce Protein A/G Magnetic Beads and incubated for 1h under mild rotation. Unbound sample was removed and beads were washed twice with lysis buffer. Bound protein was eluted by incubation with 50 ng/µl of HA peptide for 30 min. Protein concentration was normalized via Bradford assay prior the DUB activity assay.

### DUB activity assay

The DUB activity was measured using the fluorogenic substrates Ub-AMC (R&D Systems, ex 345nm, em 445nm) or Ub-Rho (LifeSensors, ex 485nm, em 535nm) in reaction buffer containing 50 mM Tris-HCl pH 7.5 and 150 mM NaCl at 30°C. When specified, 5 min pre-incubation with peroxides was carried out before the start of the reaction. For measuring DUB activity in the total cell extract, cell lysis was performed in the absence of IAA.

### Immunoblotting

Proteins were separated by standard 10-15% SDS-PAGE and transferred to PVDF membrane (ThermoFisher) for immunoblotting. The antibodies used in this work were: anti-K63 ubiquitin (1:8,000, EMD Millipore, cat. # 05-1308, clone apu3), anti-ubiquitin (1:10,000; Cell Signaling Technology, cat. #3936S), anti-uL5 (1:6,000; Cell Signaling, cat. #18163), anti-uS3 (1:6000; Cell Signaling, cat. #9538), anti-PGK1 (1:8,000; Invitrogen, cat. #22C5D8), anti-GAPDH (1:3,000, Abcam, cat. #ab9485), anti-HA (1:3,000, ThermoFisher, cat. #71-5500), anti-actin (1:5,000, Cell Signaling, cat. #4967), and anti-Rabbit IgG (1:4,000-10,000; Cytiva, cat. #NA934). Immunoblots were developed by chemiluminescence using the Amersham ECL Prime (Cytiva, cat. #RPN2232).

### ADCB sensitivity assays

Yeast cell cultures were grown into SD-Ura until log phase and diluted back to OD_600_ 0.1. Cells were mixed 1:1 with fresh medium, or medium containing 200 µg/ml of ADCB (100 µg/ml final concentration). Cells were grown into 96-well plates in triplicate and incubated for up to 96h at 30°C under agitation. Absorbance was measured at 600 nm every 15 min in a Tecan Sunrise microplate reader.

### 3D structural analysis

3D graphical images of Ubp2 structure were generated in ChimeraX (33, 34, 59) using the predicted models deposited in Alphafold2 protein database and UniProt (ID: Q01476).

### Mass spectrometry analysis

#### Sample preparation

Three independent biological replicates of WT and *ubp2Δ* cells were grown in SD complete until OD_600_ 0.4. Samples were collected at time 0-, 30- and 120-min following incubation with 0.6 mM H_2_O_2_. Cell lysis was carried out in buffer containing 50 mM ammonium bicarbonate (AmBic) pH 8, 50 mM NaCl, 1X protease inhibitor (EDTA-free), and 20 mM chloroacetamide (CIAA). Fifteen micrograms of protein were adjusted to 5% SDS in 50 mM triethylammonium bicarbonate, pH 8.5 (TEAB). Samples were reduced with 10 mM DTT for 10 min at 80°C using a Thermomixer at 1,000 rpm, alkylated with 20 mM IAA for 30 min at room temperature, then supplemented with a final concentration of 1.2% phosphoric acid and 200 µl of S-Trap (Protifi) binding buffer (90% MeOH/100 mM TEAB). Proteins were captured on the S-Trap micro device and washed 4x with 150 µl binding buffer, digested using 25 µl of 40 ng/µl sequencing-grade modified trypsin (Promega) for 1h at 47°C, and eluted using 40 µl 50 mM TEAB, followed by 40 µl of 0.2% FA, and 35 µl of 50% ACN/0.2% FA. All samples were then lyophilized to dryness and were reconstituted in 30 µl of 1% TFA/2% MeCN. A study pool QC (SPQC) was created by combining equal volumes of each sample.

#### Quantitative data-independent acquisition (DIA) LC-MS/MS Analysis

Quantitative LC/MS/MS was performed on 1 µl of each sample and replicates of an SPQC pool, using a Vanquish Neo LC coupled to a Thermo Orbitrap Astral via a Nanospray Flex ionization source. Briefly, the sample was first trapped on a Pepmap Neo Trap Cartridge and separated using on a 1.5 µm PepSep 150 µm ID x 8 cm column with a gradient of 5-12% MeCN from 0-3 min and 12-30% MeCN from 3-20 min, a flow rate of 500 nl/min with a column temperature of 45°C. The LC was interfaced to the MS using a PepSep Sprayer and stainless steel (30 µm) emitter. The MS analysis used a 240,000-resolution precursor ion (MS1) scan from 380-9080 m/z, AGC target of 500%, and maximum injection time (IT) of 50 ms, collected every 0.6 s in centroid mode. MS/MS was performed using a DIA method with default charge state of 3, precursor mass range of 380-480, 4 m/z isolation windows, AGC target of 500%, maximum IT of 6 ms, and a NCE of 28. An RF lens of 40% was used for MS1 and DIA scans.

#### Quantitative analysis of DIA data

Raw MS data was demultiplexed and converted to .htrms format using HTRMS converter and processed in Spectronaut 18 (18.4.231017.55695]; Biognosys). A spectral library was built using direct-DIA searches against a *S. cerevisiae* database, downloaded from Uniprot and appended contaminant sequences using FragPipe. Search settings included N-terminus trypsin/P specificity up to 2 missed cleavages; peptide length from 7-52 amino acids with the following modifications: fixed carbamidomethyl (Cys), oxidation (Met) and acetylation (protein N-terminus). For DIA analysis, default extraction, calibration, identification and protein inference settings were used. Peptide and protein quantification were performed at MS2 level with q-value sparse settings (precursors that met a q-value <0.01 in at least one run were included for quantification). For analysis in Spectronaut, local normalization was performed and protein abundances were calculated using the MaxLFQ algorithm (60, 61). The intensity values of 179 proteins with missing values (equal to or fewer than three) across the 18 samples were imputed using the impute.known function. This function employs the k-nearest neighbor method (k = 5) and is part of the impute R package (40). In total, 4342 proteins were included in the downstream analysis.

#### Statistical methods and visualization

Statistical tests and visualizations were conducted using R v4.0.2. Pairwise comparisons were assessed using the paired Student’s t-test, and unpaired comparisons were performed using the unpaired Student’s t-test. P-values < 0.05 were considered significant. Volcano plots were generated using the ggplot2 package (62). In Figure 7*, A-C*, proteins were color-coded based on their functions, which included antioxidant proteins, ribosomal proteins and those associated with the GCN4 regulon (63, 64). Heatmaps were created with the R package ComplexHeatmap (65), and clustering was conducted using the k-means method. GO enrichment analysis was done using DAVID functional annotation clustering (66).

## Data availability

The LC-MS/MS proteomics data (.RAW files) have been deposited to the Massive repository partnered with ProteomeXchange with the dataset identifier PXD051667. The methodology utilized to generate the proteomics dataset is described under Experimental Procedures.

## Supporting information

This article contains supporting information.

## Acknowledgments

We thank Matt Foster and the Duke Proteomics and Metabolomics Shared Resource for support with mass spectrometry data acquisition. We are indebted to Nicholas Brown and Michael Emanuele for kind donation of plasmids. We also thank Richard Brennan for kindly making SEC available. We also would like to thank the members of the Silva Lab for constructive feedback during execution of this project and preparation of the manuscript.

## Author contributions

G.M.S. conceived, supervised, and funded the study; C.M.S., B.C., D.O., N.S., N.M., V.S., and G.M.S. generated resources, performed experiments, and analyzed data. C.Y.C. performed data analysis and E.W. provided resources and supported data acquisition. C.M.S. and G.M.S. wrote the manuscript. All authors edited and contributed to its final form.

## Funding and additional information

This work was supported with funds from NIGMS R35 Award and the Chan Zuckerberg Initiative (GM137954 and SDL2022-253663, respectively, to G.M.S.). The content is solely the responsibility of the authors and does not necessarily represent the official views of the National Institutes of Health.

## Abbreviations and nomenclature

AMC: 7-amido-4-methylcoumarin (AMC)
ACN: acetonitrile
BSA: bovine serum albumin
CIAA: chloroacetamide
CHX: cycloheximide
CHP: cumene hydroperoxide
DIA: data-independent analysis
DUB: deubiquitinating enzyme
DTT: dithiothreitol
FA: formic acid
GST: glutathione-S-transferase
GFP: green fluorescent protein
H_2_O_2_: hydrogen peroxide
IP: immunoprecipitation
IAA: iodoacetamide
IPTG: isopropyl ß-D-1-thiogalactopyranoside
K48-ub: K48-linked polyubiquitin chains
K63-ub: K63-linked polyubiquitin chains
ADCB: L-Azetidine-2-carboxylic acid
LC: liquid chromatography
MS: mass spectrometry
MeCN: methyl cyanide
PBS: phosphate-buffered saline
PMSF: phenylmethylsulfonyl fluoride
PAGE: polyacrylamide gel electrophoresis
PTM: post-translational modification
ROS: reactive oxygen species
RQC: Ribosome-Associated Quality Control
RTU: redox control of translation by ubiquitin
SDS: sodium dodecyl-sulfate
t-BHP: tert-butyl hydroperoxide
TBS: tris-buffered saline
TBS-T: tris-buffered saline with 0.1% Tween 20
Ub: ubiquitin
Ub-Rho: ubiquitin-rhodamine
USP: Ubiquitin-specific processing protease
UBP: Ubiquitin-specific protease

## Supporting information for

**Figure S1.**
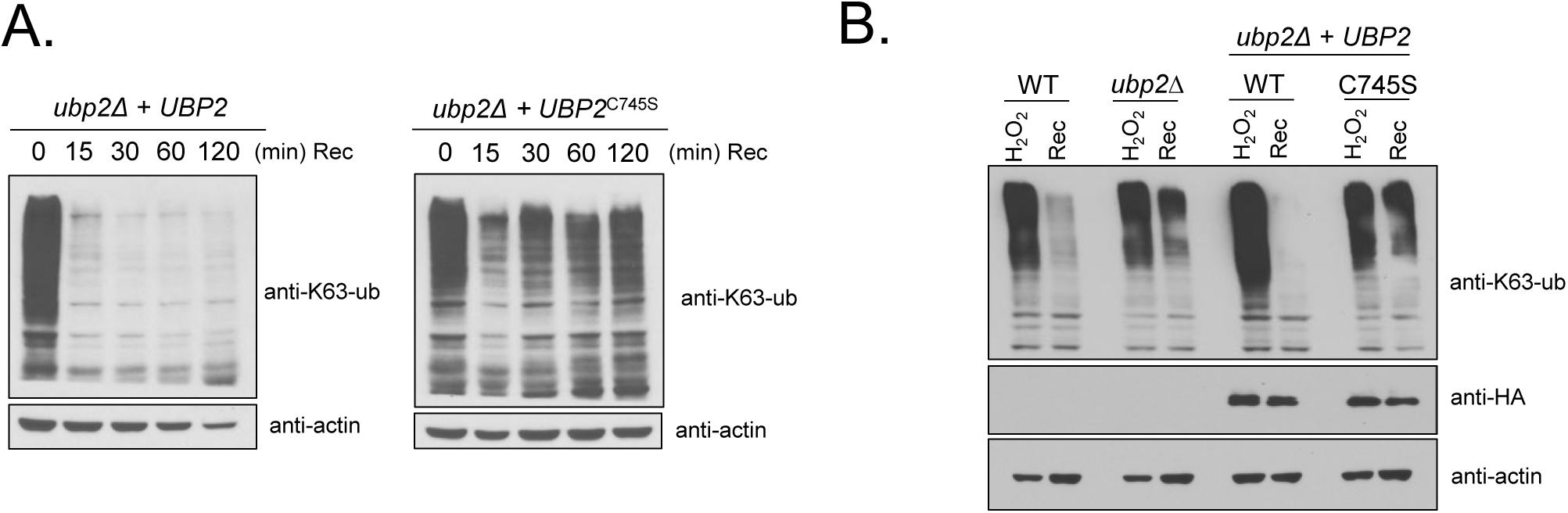
Deletion and mutation to *UBP2* impair K63-ub removal in cells. *A,* Immunoblot anti-K63-ub of lysate from cells treated in the presence or absence of 0.6 mM of H_2_O_2_ for 180 min. *B*, Total cell lysate immunoblot anti-K63-ub after 30 min of H_2_O_2_ treatment or 20 min of recovery. anti-GADPH was used as a loading control and anti-HA to detect Ubp2 levels.

**Figure S2.**
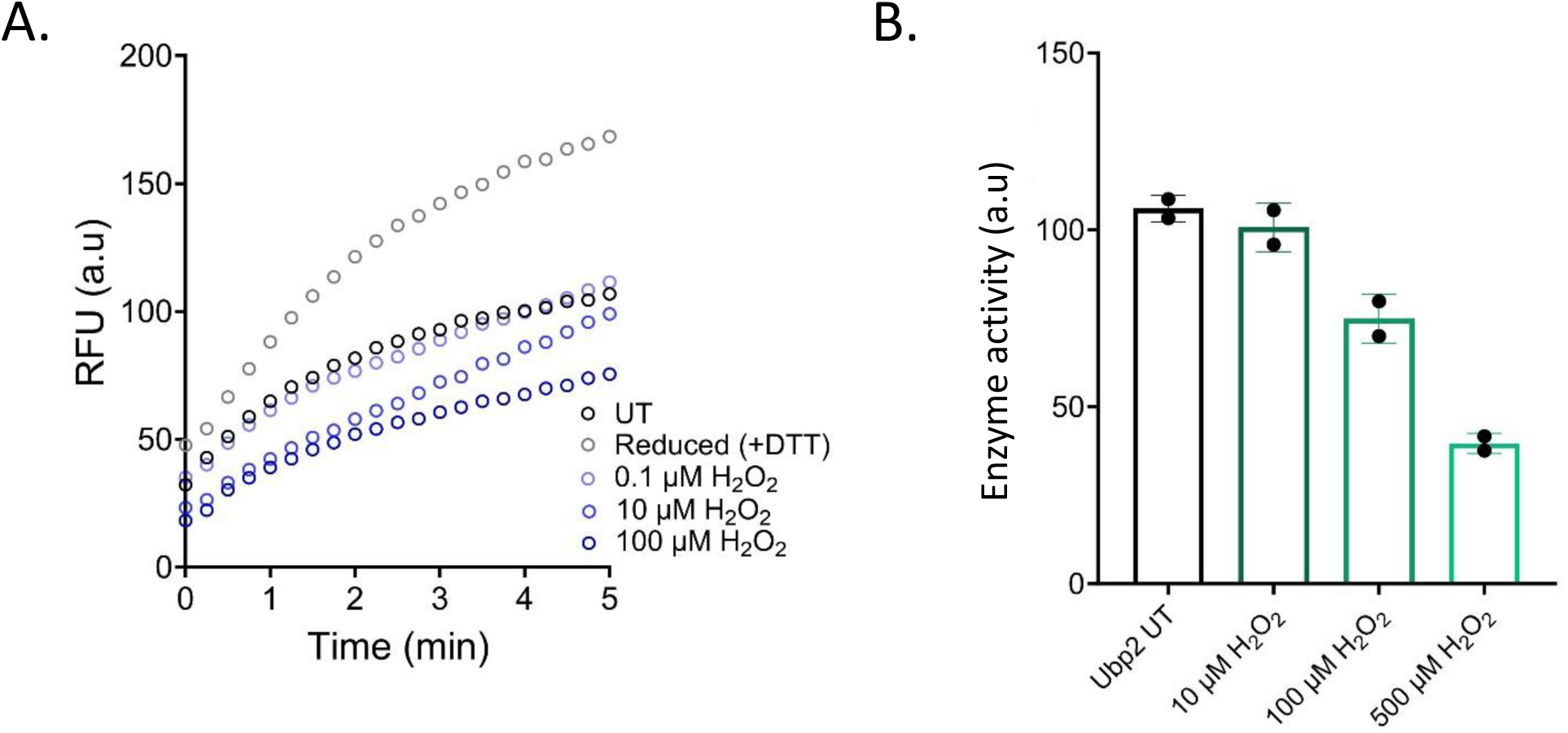
Ubp2 activity is impaired by H_2_O_2_ in a dose-dependent manner. *A*, Purified Ubp2 (100 ng) was incubated with increased concentrations of H_2_O_2_ for 5 min and DUB activity was measured with 1.5 µM of Ub-AMC. *B*, Purified Ubp2 (70 ng) was treated as above and incubated with 0.75 µM Ub-Rho. AMC fluorescence was recorded at 445 nm with excitation at 345 nm, and Rho was detected at 535 nm with excitation at 485 nm.

**Figure S3.**
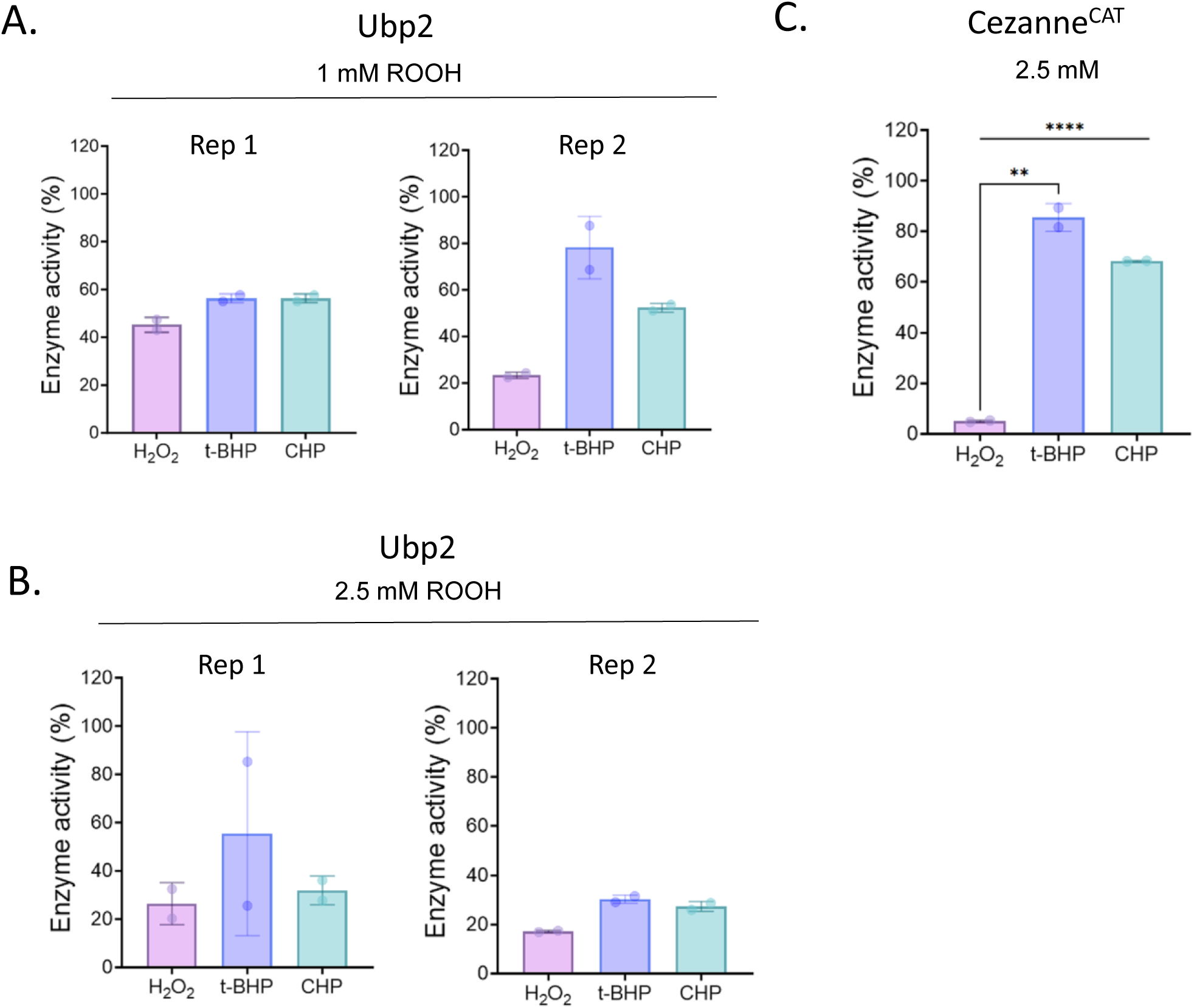
Ubp2 and Cezanne^CAT^ are sensitive to H_2_O_2 and_ organic peroxides. Purified Ubp2 (70 ng) was incubated with 1 mM (*A*) or 2.5 mM (*B*) of H_2_O_2_, t*-*BHP, and CHP for 5 min and activity was assessed by the detection of fluorescence with 0.75 µM Ub-Rho as described in Fig. S2. *C*, Purified Cezanne^CAT^ (3µg) was incubated with 2.5 mM H_2_O_2_, t*-*BHP, and CHP for 5 min and activity was assessed by the detection of fluorescence with the Ub-Rho substrate as described in Fig.S*2*. The independent purification batches (rep 1 and rep 2) are displayed. Technical replicates were averaged and Rho fluorescence was calculated as percentage of the activity of untreated Ubp2 or Cezanne^CAT^. Bar graphs show mean values ± SD for those replicates. Significance was calculated using an unpaired Student’s t-test where ** *p* <0.005, *** *p* < 0.0005.

**Figure S4.**
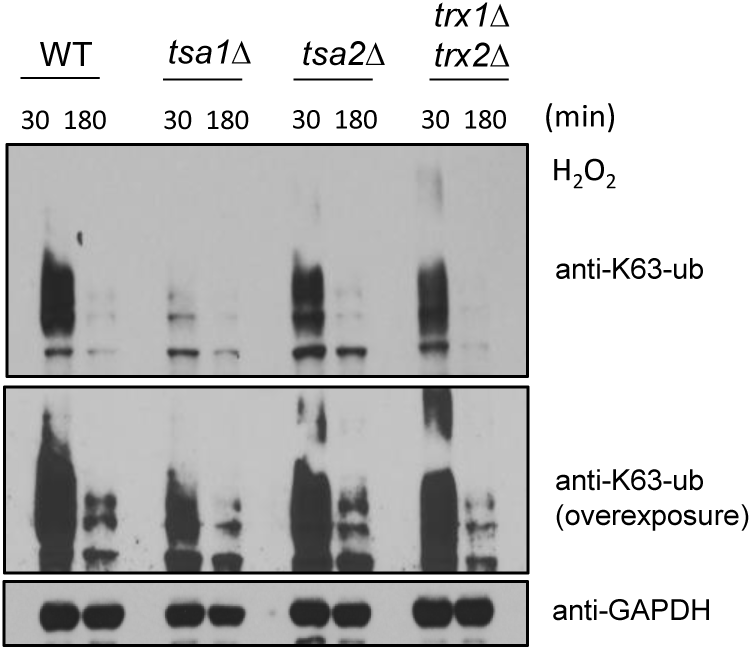
Impact of the deletion of thiol reducing system enzymes on K63-ub removal in yeast cell. Immunoblot anti-K63-ub of lysates of WT cells and cells deleted for enzymes of thiol reducing systems incubated with 0.6 mM H_2_O_2_ for the designated times. anti-GAPDH and anti-PGK1 were used as loading control.

**Figure S5.**
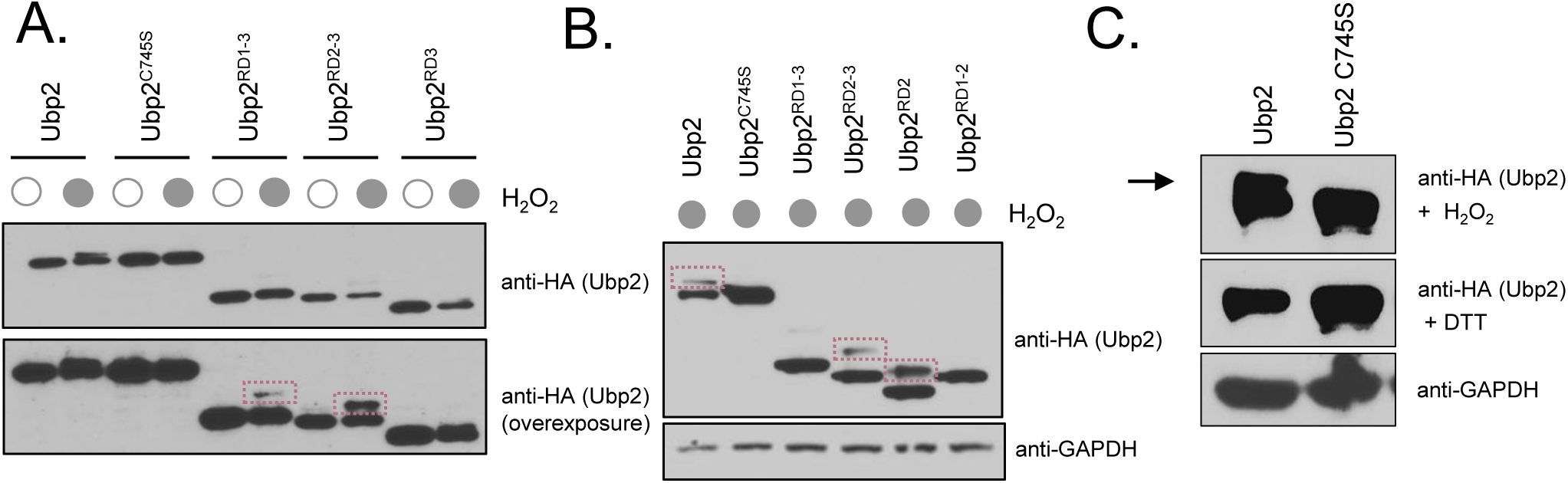
Ubp2 disulfide bonds are reversed by DTT. *A*-*C*) Immunoblot anti-HA to detect Ubp2 levels. Cells expressing HA-tagged WT Ubp2, mutants, and truncations were treated with 0.6 mM of H_2_O_2_ for 30 min and their lysates were incubated in the presence or absence of 20 mM DTT. The disulfide bond is indicated by a purple box or black arrow. Anti-GAPDH was used as loading control.

**Figure S6.**
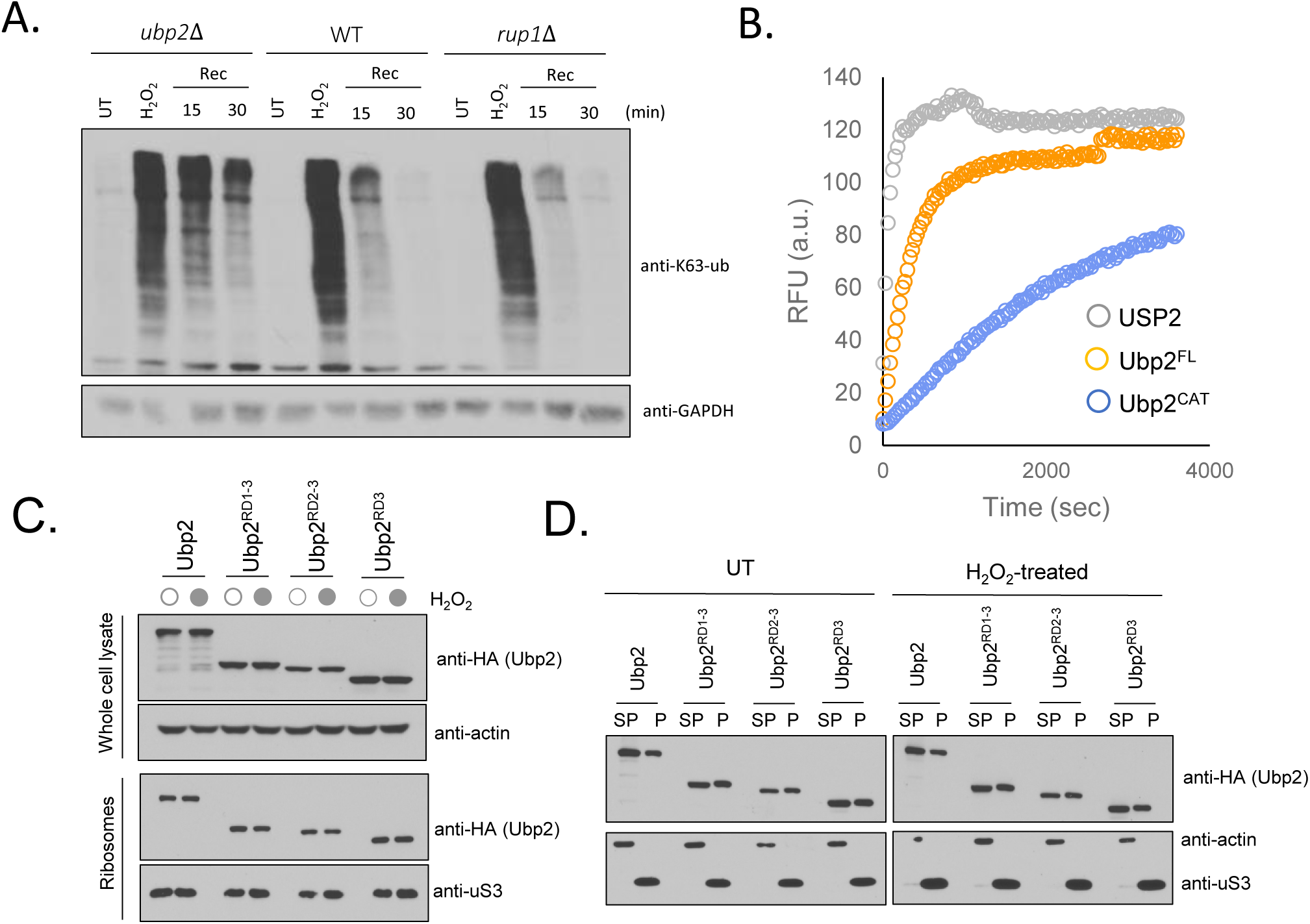
*A*, Deletion of *RUP1* does not impair Ubp2-mediated K63-ub chain reversal during stress recovery. Immunoblot anti-K63-ub of *ubp2Δ,* WT, and *rup1Δ* strains untreated or treated with 0.6 mM H_2_O_2_ followed by media swap and recovery for the designated times. anti-GAPDH was used as loading control. *B*, Ubp2^CAT^ retains partial activity *in vitro*. DUB activity for pan DUB USP2 (250 ng), Ubp2 FL (30 µg) and Ubp2^CAT^ (30 µg) was assessed *in vitro* using 1.5 μM Ub-AMC. AMC fluorescence was recorded at 445 nm with excitation at 345 nm. *C and D*, Ubp2^Nterm and^ Ubp2^RD1-3^ truncations were found associated with the ribosomal fraction in the presence or absence of H_2_O_2_. Immunoblot anti-HA of ribosomes isolated through a sucrose gradient for the HA-tagged strains shown. Supernatant (SP) and pellet (P) fractions were collected and subjected to immunoblotting. anti-uS3 was used as a loading control for isolated ribosomes, anti-actin and anti-GAPDH were used as a loading control for cell lysate.

**Figure S7.**
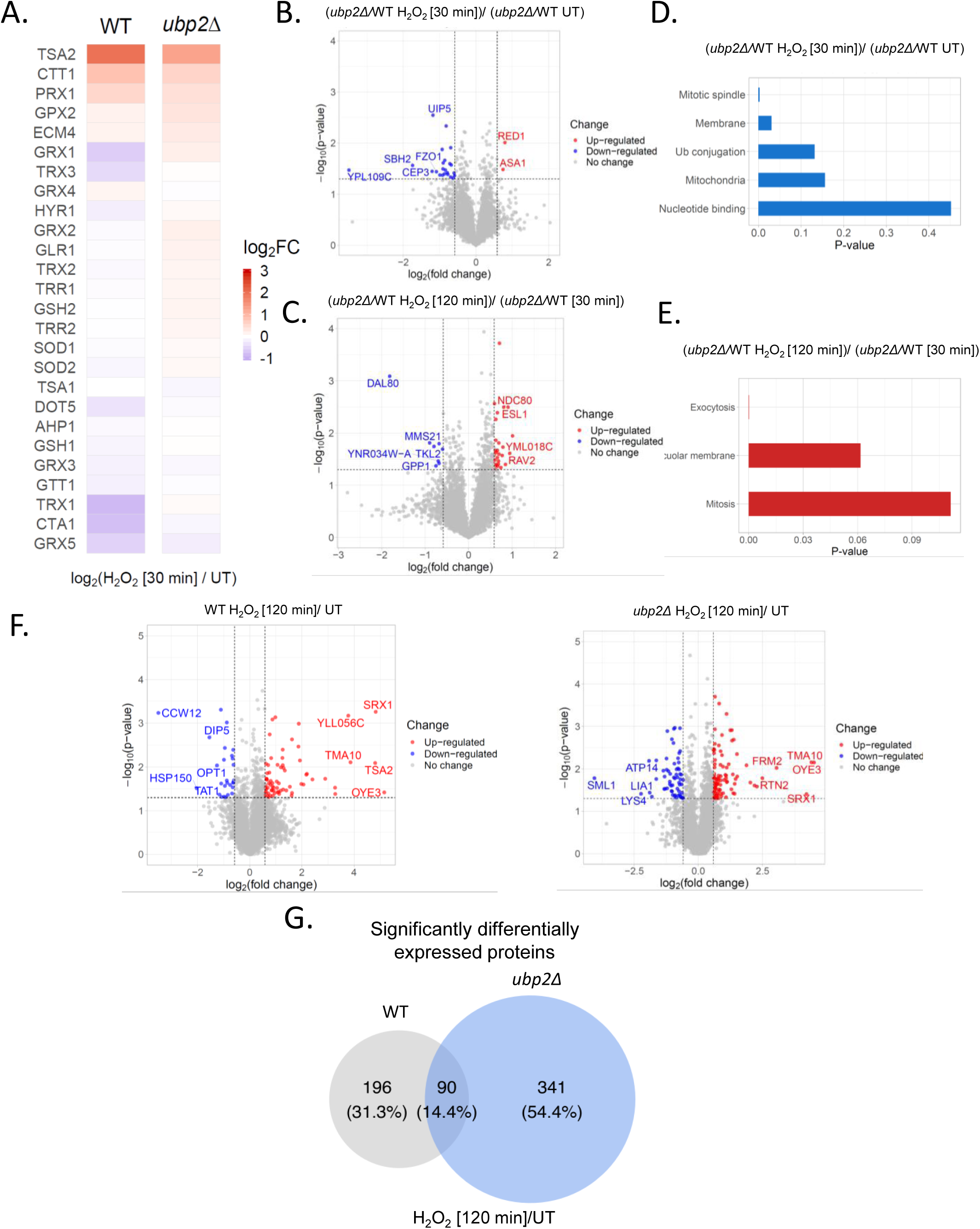
A) Heatmap of proteomics analysis showing the abundance changes for antioxidant protein levels observed in WT and *ubp2Δ* strains after 30 min of H_2_O_2_ treatment compared to the untreated (UT) condition. B and C) Volcano plot illustrating changes in protein ratios between *ubp2Δ* (30 min/0 min) vs WT (30 min/0 min) and *ubp2Δ* (120 min/30 min) vs WT (120 min/30 min) in *C*. Fold change was calculated by averaging protein ratio across replicates within each strain. *p*-values were calculated from unpaired Student’s t-tests. Significantly up-regulated proteins, with a fold change > 1.5, are colored in red, while those exhibiting significant down-regulation with a fold change <-1.5 are colored in blue. The horizontal dashed line denotes significance (*p* < 0.05), and the vertical dashed lines represent a fold change of ±1.5. D and E) GO enrichment analysis of the proteins that are down-regulated in the comparison between *ubp2Δ* (30 min/UT) and WT (30 min/UT) (*D*) and *ubp2Δ* (120 min/30 min) and WT (120 min/30 min) (*E*). *p*-values from DAVID functional annotation clustering were plotted. F) Volcano plots depicting changes in protein levels observed in the WT (*left*) and *ubp2Δ* (*right*) strains 120 min after H_2_O_2_ treatment compared to the untreated (UT). Analyses were conducted as above. *G*) Venn diagram showing proteins differentially expressed in WT and *ubp2Δ* cells between 120 min after H_2_O_2_ treatment and untreated (UT) conditions.

